# Small things matter: Lack of extra-islet beta cells in Type 1 diabetes

**DOI:** 10.1101/2025.04.11.648319

**Authors:** Kathryn Murrall, Teifion Luckett, Christiana Lekka, Christine S. Flaxman, Rebecca Wyatt, Pouria Akhbari, Irina Kusmartseva, Stephanie L. Hunter, Pia Leete, Isabel Burn, Elena Osokina, EXE-T1D consortium, James A. M. Shaw, Noel G. Morgan, Sarah J. Richardson

## Abstract

Recent 3D analyses reported abundant, small beta-cell-rich endocrine objects (EOs) in the human pancreas. Here, we assessed morphological parameters of >262,000 EOs in pancreas sections from 220 donors with or without type 1 diabetes (T1D), ranging in age and disease duration. We observe many insulin (Ins)+/glucagon (Gluc)-EOs in donors without diabetes. Their relative contribution to the total endocrine area is greatest in early life (0-2y) but reduces thereafter. Strikingly, we show the virtual absence of Ins+Gluc- EOs in individuals with T1D, where only the medium and large EOs retain beta cells. We also report a lower EO density in T1D, especially in individuals diagnosed in early life. These findings suggest that extra-islet beta cells are impacted in the development of T1D, and their early loss is a characteristic feature. This new understanding has important implications for defining beta-cell mass, which may inform future screening and treatment strategies in T1D.

**Highlights:**

- The present extensive 2D studies confirm and extend recent 3D analyses of human pancreata, demonstrating that 50% of endocrine objects (EOs) are much smaller than classical islets of Langerhans and consist predominantly of beta cells (Ins+).
- Small insulin+ EOs comprise the largest proportion of the total endocrine area in early childhood and persist throughout the life course in donors without diabetes.
- There is a shift towards larger EO size with increasing age, with the most pronounced changes in size occurring in the first few years of life.
- Small Ins+ EOs are virtually absent in individuals with type 1 diabetes, while the persisting EOs with beta cells are larger, suggesting a selective destruction of beta cells in small EOs.
- Development of type 1 diabetes, particularly at an early age, is associated with fewer larger EOs in adulthood. This implies that the lack or early destruction of small Ins+ EOs may be detrimental to the generation of larger EOs.

## Introduction

Type 1 diabetes (T1D) is an autoimmune disease in which auto-reactive immune cells selectively target pancreatic beta cells, leading to insulin insufficiency. The incidence of T1D is rising globally, with 16.4 million people predicted to be affected by 2040 (t1index.org). Increasingly recognised as a condition with equivalent rates of onset in children and adults^1,2^, T1D impacts heavily on individuals living with the condition, their families, and global healthcare systems. Hence, there is a pressing need to improve the treatment and management of the condition. Previous studies identified heterogeneity in the aetiopathology of T1D linked to age at onset, and in the clinical course of the disease, both of which are likely to impact therapeutic strategies^3–6^.

Global initiatives aimed at improving the understanding of underlying disease mechanisms in T1D recognise the need for systematic collection and collation of pancreas tissue from individuals with, or at risk of developing, T1D as well as individuals without diabetes (ND)^7,8^. The understanding that human pancreas development^9^ and T1D^10^ are not fully recapitulated in rodent models emphasises the importance of this endeavour. Several biobanks are established which facilitate the application of multi-omic 2D and 3D histological approaches to explore pancreas growth and development during the life course, however most focus on fetal or adolescent and adult tissues^11–14^. The recent establishment of the Human Neonatal Development & Early Life Pancreas Atlas (HANDEL-P) seeks to address this, but there remains a significant gap in our knowledge regarding pancreas development pre-puberty, particularly in the first two years of life.

A recent study exploited advances in 3D imaging modalities to explore volumetric endocrine mass throughout the entire pancreas of adult human donors without diabetes^15^, revealing two particularly striking findings. Firstly, 40-50% of the endocrine objects (EOs; defined as cells expressing islet hormones) are small (defined as <115 µm diameter^15^), comprising single cells or small clusters of cells which do not conform to the classical islet structure. Secondly, and in confirmation of earlier work from a small number of donors^16–20^, most small EOs are comprised exclusively of beta cells^16–20^. However, most histological studies have focussed on medium/large structures, typically referred to as pancreatic islets, leaving these small EOs underexplored. Furthermore, functional, transcriptional, and proteomic studies performed using isolated human islets do not provide any insights into the single cells and small clusters as these are lost during the isolation process^21^. Consequently, there is limited information on whether EO size and composition change during normal early-life development and with T1D. We now address these knowledge gaps by characterising the EO profile in the human pancreas throughout early development, and in T1D, using immunohistochemically labelled tissue sections from archival and contemporary pancreas biobanks.

## Results

### AI-assisted image analysis allows accurate quantification of changes in pancreas architecture and composition

Two organ donor pancreas biobanks exist where pancreas weight is available from donors collected throughout the life course: Network for Pancreatic Organ Donors with Diabetes (nPOD) and Quality in Organ Donation (QUOD-PANC). Combining these datasets confirms that the pancreas grows continuously over the first 30y of life, after which pancreas weight is relatively stable (**Fig. 1A**). When comparing fold change in pancreas weight with increasing age, the most pronounced expansion occurs within the first 2-3 years (**Fig. 1B**).

**Figure 1.** AI-assisted whole slide imaging analysis performed to investigate the development of the endocrine architecture post birth during pancreas expansion. (A) Scatter plot of pancreas weight with age post-birth with Locally Estimated Scatterplot Smoothing (LOESS) regression line in nPOD donors (n=211). (B) Fold change in pancreas weight relative to previous age bracket. (C) Whole slide image analysis methodology and data processing of slides labelled with either chromogranin A (CgA) and cytokeratin-19 (CK19) or insulin (Ins) and glucagon (Gluc). Single cells or clusters of endocrine cells are described as endocrine objects (EO). The EO area was used to estimate the number of endocrine cells present in each object, followed by EO transformation, to obtain the EO bin.

To explore pancreatic architectural changes at greater resolution, we accessed routinely labelled pancreas tissue slides from multiple archival and contemporary bioresources, covering the entire life course (Human Developmental Biology Resource: HDBR; Exeter Archival Diabetes Biobank: EADB; nPOD; Diabetes Virus Detection: DiViD; and EUnPOD). We established AI-assisted image analysis pipelines (**Fig. 1C**) to interrogate the immunolabelling of chromogranin A (CgA; to identify endocrine cells) and cytokeratin-19 (CK19), or combinations of insulin (Ins; to identify beta cells) and glucagon (Gluc; to identify alpha cells). Counterstaining of the sections with haematoxylin allowed visualisation of other pancreatic compartments (e.g. acinar, duct, blood vessels). In total, we analysed 334 whole slide scans across 220 donors, encompassing 262,832 EOs (CgA/CK19, n=45; Ins/Gluc, n=106; Ins/Gluc/CD45 n=85) in both recently labelled and archival sections. Individual donor demographics are included in **Supplementary Table 1**. The established pipelines and outputs are described in **Fig. 1C** and **Methods**. This approach yielded data derived from archival donors from whom no additional pancreas tissue is available, as well as from pancreata held in contemporary collections. These data allow assessment of changes throughout the life course (0–72y) in individuals with differing disease status (no diabetes – ND; autoantibody positive without diabetes - AAb+; type 1 diabetes - T1D).

Key pipeline outputs included: pancreas section area, total endocrine area (and proportion of total tissue area), endocrine object (EO) area (µm^2^) and count, beta and alpha cell area and proportion, and acinar area and proportion. The EO area data were transformed to generate bins as previously described^17^, representing stepwise increases in EO area as a simplified readout of continuous differences in EO size (**Fig. 1C** and **Methods**). For example, an EO bin of 0 equates to an area of 170-399 µm^2^ (representing approximately 1-3 cells), whereas a bin size of 6 equates to an area of 10880-21579 µm^2^, and a diameter of 122-180 µm, which is the approximate size of an islet equivalent (IEQ)^21^. The relationship between the EO bin, EO area and EO diameter is demonstrated in **Fig. 1C** (and **Supplementary Fig. 1A**). Visualisation of AI-generated annotations and QC comparisons of pipeline outputs on serially labelled CgA and insulin/glucagon sections confirmed that analysis pipelines were robust and reproducible (**Supplementary Fig. 1B)**. This quality-controlled pipeline allowed the assessment of pancreas architecture throughout the life course in individuals with differing disease status (no diabetes – ND; autoantibody positive without diabetes - AAb+; type 1 diabetes - T1D).

### Endocrine object density and area decrease during the first ten years of life in healthy donors, and the endocrine object profile shifts towards larger islets with age

Using donor tissue immunolabelled for CgA & CK19 (n=45; HDBR n=10, EADB n=35), we initially quantified total EO density (**Fig. 2A**) and percentage endocrine area (**Fig. 2B**) in ND donors. A reduction in both EO density and proportional endocrine area was apparent over the first 10 years of life and was particularly prominent during the first two years, coinciding with marked acinar expansion (**Fig. 2C** and **Supplementary Fig. 2A**).

**Figure 2.** The endocrine mass in individuals without diabetes is contained within increasingly larger endocrine objects (EO) with age, as the pancreas matures post-birth, which coincides with a reduction in EO density. **(A)** Changes in EO density with age in pancreata with No Diabetes (ND), immunolabelled with CgA/CK19 (n=45) or Ins/Gluc (n=62). **(B)** Changes in endocrine area (%) with age. **(C)** Representative images of EO clusters in ND neonatal, 3y and 10y pancreata immunolabelled with Ins/Gluc. **(D)** Proportion of total EO count and area in EO bins (0-9) in CgA/CK19 labelled ND pancreata. **(E)** Proportion of endocrine area in small EOs (bin 0-3) and large EOs (bin 7-9) across selected age brackets in CgA/CK19 labelled pancreata. **(F)** Proportion of total EO count and area in each EO bin (0-9) from Ins/Gluc labelled ND pancreata. **(G)** Proportion of endocrine area in small EOs (bin 0-3) and large EOs (bin 7-9) across selected age brackets in Ins/Gluc labelled pancreata. **(H)** Representative spatial plots of EO distribution and size in ND pancreata at birth (0y), 9y and 24y of age. Colour and size of points indicate EO bin size and class respectively. Key not to scale. Data are presented as mean ±95% CI. Kruskal-Wallis test followed by Tukey’s *post hoc* test was performed to calculate *p* values. *p <0.05, **p <0.01, ***p<0.001).

Across the 45 donor sections assessed, a total of 42,793 EO were identified. Using the EO binning strategy, we assessed the frequency EOs in each sized bin and the contribution of each bin to the overall endocrine area across different donor ages (fetal, 0-1y, 2-6y, 7-12y, 13-18y). There was a clear shift towards a greater frequency of larger EOs with age, and in the proportion of the total endocrine area allocated within larger EO bins (**Fig. 2D** and **E** and **Supplementary Fig. 2B**). The proportion of endocrine area in medium-sized bins (4-6) did not change significantly post-birth (**Supplementary Fig. 2C**). Strikingly, most (76.3-98.6%) of the EOs were small and assigned to bins 0-3 across all ages (**Fig. 2E** and **Supplementary Table 2)**. Nevertheless, the EOs within bins 0-3 contributed only a small proportion of the total endocrine area (15.9-36.7%); this was especially evident during the postnatal period but reduced further with age. The shift in EO profile with age was confirmed in a second batch of donors (n=62; nPOD = 36; EADB = 20; EUnPOD= 6) aged 0-72 years, where beta and alpha cells were visualised using insulin and glucagon immunolabelling (**Fig. 2F** and **G** and **Supplementary Fig. 2D-F**). A total of 243,326 EOs were identified. Donors were split into age categories (0-1y, 2-6y, 7-12y, 13-17y), and an additional group of ≥18y donors was included. The EO profiles in age-matched donors from contemporary organ donor (nPOD, EUnPOD), live donor (DiViD) and archival biobanks (EADB, Seattle) were concordant (**Supplementary Fig. 1C**). Assessment of density in each of the EO bins revealed a reduction with age; the greatest density of small-medium EOs (bins 0-6) was observed in the very youngest donors (0-1y) (**Supplementary Fig. 2D**).

Examination of Feret minimum and maximum diameters indicated that a standard IEQ, with a minimum diameter of at least 50 µm, would be grouped into EO bin 4 or above (**Supplementary Fig. 1A**). Therefore, all EOs allocated to bins 0-3 are likely to be excluded from the preparations of islets obtained during pancreas digestion and islet isolation^21^. This represents a large proportion of the total EOs (73-87%) and the proportional endocrine area (19-49%) in a given individual (**Fig. 2F**).

Age-dependent architectural changes are visually apparent in representative spatial plots of EOs, which are proportionally sized and coloured according to bin size (**Fig. 2H** and **Supplementary Fig. 3A**). These plots illustrate the pronounced increase in EO size and the corresponding reduction in EO density with age.

### Hormone immunolabelling-based class splitting of EOs reveals that most small clusters comprise exclusively beta cells

In their recently published 3D analysis of the whole human adult pancreas, Lehrstrand *et al*.^22^ separated EOs based on the presence of insulin alone (beta cells only; Ins+Gluc-), insulin and glucagon (beta and alpha cells; Ins+Gluc+) and glucagon alone (alpha cells only; Ins-Gluc+). In the present analysis, the insulin and glucagon positive areas within each distinct EO from individuals with (T1D; n=114 donors; 88,617 EOs) and without diabetes (ND; n=62; 109,936 EOs) were identified. EOs were segregated into three classes based on whether the total insulin/glucagon-labelled area exceeded 40 µm^2^. Comparing the proportion of the total EO count across these three classes in ND donors revealed that Ins+Gluc- EOs were most frequent (Ins+ Gluc- 54%±3.3%.; Ins+Gluc+ 37.1±2.5%; Ins-Gluc+ 8.9±0.8%; **Fig. 3A**). However, when comparing the contribution of each class to the total EO area, the Ins+Gluc- EOs represented a significantly smaller proportion of area (p<0.001) than the Ins+Gluc+ EOs (Ins+ Gluc- 17.2±2.5%; Ins+Gluc+ 81.2±2.6%; Ins-Gluc+ 1.7±0.5%; **Fig. 3A**). This implies that the Ins+Gluc- EOs, although large in number, are smaller in aggregate area; in accordance with observations from 3D imaging studies^15^. Binning Ins+ Gluc- EOs by size, shows that 96% are small and allocated within bins 0-3 (**Fig. 3B** and **Supplementary Fig. 3B and 4A)**. Similarly, Ins-Gluc+ EOs are also small, with 99.2% contained within bins 0-3. In, contrast Ins+Gluc+ EOs are predominantly large, with bins 4-8 comprising 70.3% of the total endocrine area (**Fig. 3C**). Quantification of the %Ins+ and Gluc+ areas confirmed that most of the beta cell and alpha cell area is attributed to EOs in bins 3-7; representing larger sized structures (**Fig. 3D**).

**Figure 3.** **Beta cells contained within larger EOs, which are more prevalent in older individuals, are more likely to persist in type 1 diabetes. (A)** Proportion of total EO count and area in each of the three EO classes (Ins+Gluc-; Ins+Gluc+; Ins-Gluc+) in ND and T1D pancreata. Mann-Whitney U test with Benjamini-Hochberg correction performed to calculate adjusted *p* values. **(B)** EO density in bins separated by class for ND and T1D pancreata. Type II ANOVA followed by Tukey *post hoc* comparing ND-T1D for each bin was performed to calculate *p* values. **(C)** Proportion of endocrine area in each EO bin separated by class. **(D)** % Insulin and glucagon-labelled area as a proportion of the tissue area in each EO bin. **(E)** Proportion of EO area (%) in each bin separated by class, for ND and T1D (duration <2y) pancreata, grouped by age. **(F)** Representative spatial plots of ND and T1D (duration <2y) pancreata, from neonates, children and adults. Each EO is represented by a single point whose size and colour correspond to EO bin and class, respectively. **(G)** Proportion of EO area contained within Ins+Gluc+ and Ins-Gluc+ EOs in ND and T1D pancreata, from donors aged 1-12y and ≥13y. Data are presented as mean ±95% CI. *p <0.05, **p <0.01, ***p<0.001).

### Beta cells as single cells or in small clusters are virtually absent in type 1 diabetes

A comparable analysis performed in donors with T1D (**Fig. 3A**) revealed a smaller total proportion of Ins+Gluc- EOs (ND 54±3.3% v T1D 5.8±2.5%; p<0.001) and their total area (ND 17.2±2.5% v T1D 1.6±0.9%, p<0.001). The Ins+Gluc+ EOs also made up a smaller proportion of total EOs in individuals with T1D (ND – 37.1±2.5%, T1D – 12.6±2.5%, p<0.001); their corresponding area was also smaller (ND – 81.2±2.6%, T1D – 36.4±5.6%, p<0.001). The inverse was observed with Ins-Gluc+ EO count (ND – 8.9±1.6%, T1D – 81.6±2.1%, p<0.001) and area (ND – 1.7±0.5%, T1D – 62±5.8%, p<0.001), which were greater in T1D than ND.

Assessment of the EO bins in individuals with T1D confirmed that the smallest Ins+Gluc+ bins were most impacted, with significantly lower EO density, % endocrine area and % of total EO count in bins 0-4 compared to ND individuals (**Fig. 3B** and **C** and **Supplementary Fig. 4A**; p<0.001). Intriguingly, the residual Ins+Gluc+ EOs in the T1D donors exhibited a rightward shift compared to ND, suggesting that the larger Ins+Gluc+ EOs were preferentially preserved in T1D. The contribution of Ins-Gluc+ EOs to endocrine area in bins 1-7 was significantly greater in the T1D donors compared to ND (p<0.001). Most of the Ins-Gluc+ EO area was contained within bins 3-8 (50.8% proportion of total endocrine area and 81.8% of total Ins-Gluc+ area), confirming residual EOs retain alpha cells in the absence of beta cells. Furthermore, a significantly greater number of Ins-Gluc+ EOs in smaller bins implies more single alpha cells and small clusters of alpha cells, which are not frequently observed in ND donors. Assessment of total Ins+ and Gluc+ endocrine areas in T1D confirmed the significant loss of Ins+ cells across the first seven EO bins. There was a significant increase in Gluc+ area relative to acinar area in bins 2-6 (**Fig. 3D**). It is not clear whether this represents an increase in alpha cell number/area (an increase in cycling alpha cells was reported recently in individuals with diabetes^23,24^), a reduction in acinar area, or an increase in beta cell transdifferentiation^25^. Consistent with the lower frequency of the small beta cell only EOs and the trend towards larger residual Ins+Gluc+ EOs in donors with T1D, the residual Ins+ area was predominantly confined to EO bins 4-9, suggesting that beta cells within smaller EOs may be more susceptible to immune-mediated destruction. Assessment of further morphological measurements of EOs in individuals with T1D confirmed a reduction in circularity and solidity with increasing EO bin size, also seen in ND donors, although this was more prominent in bins 3-7 in individuals with T1D (**Supplementary Fig. 4B and C)**.

### Age at diagnosis of type 1 diabetes correlates with endocrine object bins and classes

We previously characterised two pancreatic endotypes of T1D which are closely correlated with age at clinical diagnosis^3,4^. Diagnosis under the age of 7 years is associated with the most intense immune infiltration and a reduced frequency of residual insulin-containing islets at onset (type 1 diabetes endotype 1: T1DE1), when compared with individuals diagnosed later in life (≥13y; type 1 diabetes endotype 2: T1DE2)^3,4^. To explore the impact of age at T1D diagnosis on EOs, we assessed EO parameters in donors with a short duration of T1D (<2y; n=70) and compared the outcomes with age-matched ND donors. Donors were divided into five age categories (0-1y, 2-6y, 7-12y, 13-17y and ≥18y) to enable interrogation of both pancreatic architectural changes and the impact of disease. A clear age-dependent gradient in EO parameters was observed, where the individuals diagnosed at younger ages exhibited an increasingly and incrementally severe loss of Ins+Gluc+ EOs, measured by proportion of total endocrine area (**Fig. 3E**) and proportion of total EO count (**Supplementary Fig. 4D**). The virtual absence of Ins+Gluc- EOs and preservation of the larger Ins+Gluc+ EOs was apparent in donors aged 2-6y at diagnosis, when compared with their age-matched controls. Spatial plots visually represent these observed changes and highlight the lobularity of disease, which is frequently reported (**Fig. 3F**). These findings support the conclusion that beta cell loss is more profound in individuals who are diagnosed with T1D at an early age, compared with those who are older at onset.

Assessment of the proportion of the endocrine area accounted for by Ins+Gluc+ and Ins-Gluc+ classes in individuals with short-duration disease (<2y) and in ND donors split by donor age (1-13y versus ≥13y; **Fig. 3G**) strengthens these findings. Comparison of the median ages of the ND and T1D groups (ND 1-12y: 6y, T1D 1-12y: 7y and ND ≥13y: 31y, T1D ≥13y: 22y), suggests the differences observed are due to diabetes status rather than pancreas maturity. These data demonstrate that Ins+Gluc+ EOs account for a substantially smaller proportion of the total endocrine area, and the residual Ins+Gluc- EOs a larger proportion, in individuals diagnosed <13y (mainly T1DE1) compared to age-matched controls (%Ins+Gluc+ area; ND 1-12y: 82%, T1D 1-12y: 31.6%; ND ≥13y: 82.4%, T1D ≥13y: 57.6%). In contrast, the reduction in Ins+Gluc+ EO area and the EO bin size alterations are less apparent in donors diagnosed ≥13y (T1DE2). Furthermore, when comparing the Ins+Gluc+ and Ins+Gluc- proportional areas in children diagnosed between 1-13y, a significant reduction in EO bin size is evident, which may reflect the comprehensive destruction of beta cells within these EOs. The shift in EO bin area between Ins+ EOs and Ins-EOs is much less apparent in donors diagnosed ≥13y. This may be explained by the initiation of dedifferentiation or transdifferentiation events and/or comparatively reduced endocrine cell loss from affected islets.

### Endocrine object parameters are not impacted in single autoantibody positive donors without diabetes

We demonstrated changes in EO area, density, and morphology consistent with age and progression of T1D. However, to determine whether T1D-associated alterations are due to disease, rather than a predisposing factor such as a developmental impairment, we assessed relevant parameters in 15 AAb+ donors. Initially, donors were classified according to the number of autoantibodies detected (**Fig. 4A**) and their age (2-6y; ≥18y).

**Figure 4.** The presence of single or multiple autoantibodies in most individuals is not associated with a loss in small Ins-Gluc+ EOs. **(A)** EO density in bins separated by class in age-matched ND, single autoantibody positive, double autoantibody positive, and T1D pancreata. Type II ANOVA with Dunnett’s *post hoc* test comparing single and double autoantibody positive to control ND pancreata was performed to calculate *p* values for each EO bin comparison. **(B)** The sum of small Ins+Gluc- EO density (bin 0-3). Data point colour indicates detected autoantibodies. Kruskal Wallis test followed by Tukey’s *post hoc* test was performed to calculate *p* values. **(C)** Spatial plots comparing age-matched ND and single autoantibody positive ND pancreata. Data are presented as mean ± 95% CI. *p <0.05, **p <0.01, ***p<0.001.

There was no difference in the density of small EOs between AAb+ individuals and age-matched ND donors in the 2-6y group, and no difference between donors with one AAb and ND donors in the ≥18y group (**Fig. 4A** and **B**) even if the donors scored highly for T1D-genetic risk score (T1D-GRS)^26^ (**Supplementary Table 3**). Spatial visualisation of AAb+ and ND donor pancreas sections confirms similarity (**Fig. 4C)**.

### Endocrine object parameters are impacted in autoantibody positive donors without diabetes that have additional features associated with progression towards clinical T1D

Donors with two autoantibodies showed a trend towards fewer Ins+Gluc- EOs in bins 1-3 (bin 1, p<0.05) and a significantly smaller proportion of total EO count in bins 1-2 (**Fig. 4A** and **Supplementary Fig. 5A)**. Furthermore, greater density, proportion of EO count and endocrine area of small Ins-Gluc+ EOs was observed in ND donors with two autoantibodies (**Supplementary Fig. 5B and C**). However, this shift in the density and proportion of small EOs did not coincide with a change in the % endocrine area. Emerging evidence suggests that AAb+ individuals immunopositive for insulinoma associated protein 2 (IA2A) are at greater risk of progressing towards clinical diabetes^27^. We observed that donors with IA2A+ and one other autoantibody trended towards a lower density of Ins+Gluc- EOs and a higher density of Ins-Gluc+ EOs (**Fig. 4B)**. Therefore, donors were subdivided according to the presence or absence of IA2A, versus other single AAbs, or two other AAbs for further analysis. The donors with IA2A+ and one other autoantibody, all aged 18y (n=3), had significantly fewer Ins+Gluc- EOs in bins 0 and 2 as a proportion of total EO count, and significantly more Ins-Gluc+ EOs in bins 0-1, but this did not significantly impact overall endocrine area. (**Fig. 5A** and Supplementary Fig. 5D-F).

**Figure 5.** The presence of IA2A, high T1D-GRS and histological features of diabetes progression is associated with a reduction in small Ins+Gluc- EOs and an increase in small Ins-Gluc+ EOs. **(A)** EO density in each bin separated by class in age-matched donor pancreata, grouped by presence of IA2A and numbers of autoantibodies. Type II ANOVA with Dunnett’s *post hoc* test comparing single and double autoantibody positive donors to control ND pancreata was performed to calculate *p* values for each bin. **(B** and **C)** EO density in bins separated by class **(B)**, and spatial plots **(C)** comparing IA2A+ donors to age-matched ND and T1D pancreata. Data are presented as mean ± 95% CI. *p <0.05, **p <0.01, ***p<0.001.

Islet autoantibody positivity alone is not sufficient to predict whether an individual will progress to overt clinical disease. Indeed, most donors with islet autoantibodies, particularly a single AAb, do not^28^. The T1D staging criteria predict progression using the presence of multiple AAbs, combined with measures of glucose control^29,30^. However, as accurate measures of glucose control are not available in the present study, we sought to identify donors in our cohort with other histological indicators of progression, including insulitis, hyperexpression of HLA Class I^31–33^, and a higher T1D-GRS^26^. We identified three donors with previously reported evidence of HLA-I hyperexpression and insulitis: nPOD 6267, 6167 and 6197, and elevated T1D-GRS1 scores (**Supplementary Table 3**). All were immunopositive for IA2A and either glutamic acid decarboxylase (GADA; nPOD 6267 and 6197) or zinc transporter 8 (ZnT8; (nPOD 6167). Donors 6267 and 6167 had fewer small Ins+Gluc- EOs, and a smaller proportion of insulin labelling:acinar area compared to age-matched ND donors (**Fig. 5B** and **C**). By contrast, the endocrine profile of 6197 resembled the ND profile.

In summary, in most AAb+ donors with single or multiple autoantibodies, even those with a high T1D-GRS, we observe no difference in EO distribution when compared with age-matched ND donors, implying that there is not a defect in small EO development in at-risk individuals. However, in donors with a high T1D-GRS, IA2A positivity and other histological features of T1D, some exhibit alterations in EO parameters that resemble changes observed in individuals with a short T1D duration, supporting alterations associated with active pancreatic autoimmunity.

### There is a lower density of large islets in individuals diagnosed with type 1 diabetes at an early age (<13y)

Throughout these analyses, we identified multiple small beta cell-only EOs present at a high frequency and density in young, healthy individuals. We show that these are virtually absent in individuals recently diagnosed with T1D, particularly those with a young onset of disease. This suggests that the smaller EOs may be more sensitive to immune-mediated destruction. We also demonstrate a shift in EO size towards larger EOs during early post-natal development in individuals without diabetes, indicating that the small EOs may mature into larger EOs with age. It is well established from natural history studies that individuals who develop islet AAbs at a very young age (<2y) are at greater risk of progression to clinical diabetes than those who develop AAb later in childhood^34–36^. Therefore, we hypothesise that earlier initiation of islet autoimmunity, when the developing pancreas has a high density of small EOs, results in faster and more comprehensive destruction of beta cell mass, which may impact the development of larger EOs, if small EOs are a precursor to their formation. To examine this, we assessed total EO bin density in adult individuals with T1D with a longer duration of disease (≥2y), diagnosed either <13y (mainly T1DE1) or ≥13y (T1DE2) based on our previously defined T1D pancreatic endotypes (**Fig. 6A**). Medium and large EO density (bins 3-8) was significantly less in individuals diagnosed <13y compared to those diagnosed ≥13y. This is visually apparent when examining spatial plots from representative donors (**Fig. 6B** and **Supplementary Fig. 6A-B**). These data suggest that clinical onset of T1D at an early age is detrimental to the formation of larger islets, resulting in a lower density of medium-large EOs that persists into adulthood.

**Figure 6.** Clinical onset of type 1 diabetes <13y results in more efficient destruction of the beta cell mass after 2y duration and interrupts the pathway for better protected large islets to form. (A and. **B)** Endocrine object density in bins **(A)** and representative spatial plots **(B)** of adult ND, T1D Endotype 1 (T1DE1, diagnosed <13y) and T1D Endotype 2 (T1DE1; diagnosed ≥13y) pancreata with a disease duration of ≥2y. Mann-Whitney U test and Benjamini-Hochberg procedure were used to calculate adjusted *p* values. **(C-E)** Proportion of EO count, **(C)** area, **(D)** and Ins+ or Gluc+ area **(E)** in each bin separated by class, for age-matched ND, T1DE1 (duration ≥2y) and T1DE2 (duration ≥2y) pancreata. Type II ANOVA with Tukey’s *post hoc* test comparing T1DE1 to T1DE2 for each bin, separated by class was performed to calculate *p* values. **(F)** Sum of % Ins+ area contained within medium and large EO bins (4-9). Outliers are labelled by donor ID. Mann-Whitney U test was performed to calculate *p* values. Data are presented as mean ± 95% CI. *p <0.05, **p <0.01, ***p<0.001.

### Residual Ins+ area is greater in individuals who developed clinical T1D at a later age (≥13y) and this is confined to larger EOs

Previously, clinical onset at an early age (<13y) was associated with marked loss of beta cell mass^3,4,37^. Here, we explored the contribution of each of the EO classes to the total EO count and endocrine area in age-matched individuals with a disease duration of ≥2y. By matching the donors according to age, we account for changes in acinar expansion and the increase in EO size with age. In both groups assessed (13-17y and ≥18y), we observed a trend towards greater proportions of Ins+Gluc+ EOs in bins 6-8 in T1DE2 donors, with a later clinical onset, compared to T1DE1 donors (**Fig. 6C-E**). Examination of the total Ins+ area showed that those diagnosed ≥13y trended towards greater Ins+ area in the larger EO bins than those diagnosed <13y (**Fig. 6E**). Plotting the cumulative Ins+ area in bins 4-9 shows that little beta cell mass remains in T1DE1 donors (**Fig. 6F**). In contrast, most T1DE2 donors showed greater preservation of beta cell mass in medium-large EOs, though some had equivalent beta cell mass to the T1DE1 donors, indicating greater heterogeneity in the T1DE2 donor group (**Supplementary Fig. 6 and 7**). Overall, these findings suggest that the larger EOs in older individuals are more resistant to immune attack as they retain more beta cell mass.

## Discussion

The recent 3D visualisation of an entire human pancreas^15^ emphasised the presence of small hormone-positive cell clusters located remotely from the more classically defined islets of Langerhans. The striking number of these small“endocrine objects” was largely, although not totally^16–20^, unappreciated in earlier studies. This may reflect the heavy reliance on 2D analysis of pancreatic tissue, in which clusters of cells cannot be readily assigned as “extra-islet” structures due to the topography of the tissue. By contrast, mesoscopic optical 3D imaging approaches provide unequivocal evidence that as many as 50% of the endocrine objects are smaller than classical islets and consist primarily of Ins+ beta cells, with few or no Gluc+ alpha cells^15^. The present study builds on these newer 3D^15,38^ and early 2D^16–20^ studies by applying an innovative analysis pipeline to define the endocrine cell topography in 2D pancreas sections from >200 donors across a wide range of ages, held in multiple biobanks. We confirm that among individuals without diabetes, 40-60% of total EOs exist as small clusters of cells predominantly comprising beta cells. We further demonstrate that the relative endocrine area (as a proportion of total pancreas area) and the endocrine object density each reduce consistently in the first few years of life, with endocrine area stabilising at 1-2% of total pancreas area by the early teenage years. Both an earlier^20^ and a very recent study^39^ using different pancreas biobanks reached similar conclusions.

Post-mortem studies of pancreas weight at increasing ages support the notion that the pancreas expands until the age of ∼30y^40–42^. Maintenance of endocrine area at 1-2% of the pancreas beyond the teenage years implies that endocrine cell expansion continues in some form over at least the first three decades of life. Debate as to whether this sustained increase in endocrine mass, especially beyond the age of 10y, is derived from endocrine proliferation or neogenesis is ongoing^20,43–47^. Whilst the present study cannot provide a resolution, applying our methods in future studies focussed on the proliferation status of the smaller EOs and/or their proximity to ductal structures, as potential sites of endocrine neogenesis, should provide valuable new insights.

Our analysis of thousands of pancreatic EOs throughout early life and into adulthood firmly demonstrates that the proportion of the EO area allocated within the smallest EO bins is greatest in early childhood and that a pronounced shift towards larger EOs occurs with increasing age. This change coincides with a period of rapid pancreatic growth and extensive architectural changes largely overlooked in earlier work. The lack of a thorough assessment of these changes, and the paucity of information about the likely functional significance of the small EOs given their loss during islet isolation, leaves many questions outstanding. These include the conundrum that these small EOs may be distinct entities with specific functions that differ from those attributed to a typical islet and that their functionality and topography may change throughout the life course. It is tempting to speculate that the shift towards larger EOs with age is evidence of a neogenetic pipeline culminating in larger EO (islet) generation, with EOs at different stages of this developmental process present within any particular individual^48–50^. Given that studies of pancreas tissue and isolated islets demonstrate enhanced insulin secretion, greater insulin content, a higher proportion of beta cells, and a greater percentage of beta cells in direct contact with blood vessels in smaller islets^51^, it is likely that EO size and composition are functionally important.

Arguably, this study’s most striking and significant observation is the virtual absence of small EOs in the pancreas of donors with T1D. This implies that the smaller, beta-cell-rich objects may be particularly susceptible to autoimmune-mediated destruction. These observations align with 3D imaging studies in the non-obese, autoimmune mouse model, where smaller islets are also least resistant to immune attack^52^. In contrast, in the streptozotocin (STZ)-induced model of diabetes, where beta cell toxicity rather than autoimmunity causes beta cell loss, the small EOs are preserved, with the larger islets impacted^53^. This suggests a functional difference between the beta cells in small clusters vs classical islets, at least in mice, since STZ requires the expression of GLUT2 for its uptake, which may be lacking in extra-islet beta cells. However, the impact of STZ on endothelium and its consequential impact on small EOs cannot be ruled out^54,55^. Furthermore, despite our inability to examine in detail immune infiltration around small EOs in human T1D pancreas, due to their virtual absence, the evidence of elevated acinar inflammation in individuals with T1D, and the presence of islet-reactive CD8+ T cells in the exocrine compartment supports the presence of an immune component capable of targeting beta cells in the extra-islet environment^56–58^. This could reflect remnants of active beta-cell autoimmunity and aligns with the persistence of Ins-Gluc+ small EOs. Consistent with this hypothesis, residual beta cells were predominantly confined to larger EOs, particularly in donors clinically diagnosed at the youngest ages. This suggests that beta cells contained within the large EOs may be initially protected from autoimmune-mediated destruction and that smaller EOs are lost early in the disease process. Factors such as the presence of an intact islet basement membrane (reducing accessibility of immune cells); the localisation and number of protective accessory cells (e.g. pericytes, mesenchymal stromal cells); and/or the beta cells mounting an active repulsive response could provide an explanation^59–63^.

We also observed a lower EO density in individuals with T1D compared to age-matched ND donors. This aligns with an earlier study undertaken in a limited number of T1D donors diagnosed in adulthood^64^. This lower density was particularly apparent for medium-large EOs in children who developed diabetes at <13y vs age-matched ND individuals. This implies that the selective targeting of small EOs early in the autoimmune process could negatively impact the formation of larger EOs during childhood, thereby resulting in a reduced density of all EOs in later life. Hence we propose that, particularly in young children, T1D may be a disease of beta cell loss (driven by a more aggressive immune-mediated destruction^3,4^) coupled with a net initial beta cell deficit, resulting from a failure to establish larger endocrine objects in sufficient quantities. In individuals diagnosed in their teens and beyond, beta cells are preferentially preserved in the medium and large EOs for extended periods post-diagnosis, perhaps as these are formed prior to the initiation of the autoimmune attack. These findings align with clinical observations showing that individuals diagnosed beyond their earliest years are more likely to retain the ability to make at least some endogenous insulin (assessed by C-peptide microsecretion)^37,65^, and exhibit less florid insulitis^4,66^. Nevertheless, cross-sectional studies measuring C-peptide decline with increasing disease duration support a continuous loss of residual functional beta cells (at least for the first 7 years)^65^. Notably, when comparing the size of the Ins- and Ins+ EOs, the shift in bin size with age was less pronounced in individuals diagnosed ≥13y than in individuals diagnosed <13y. This corresponds with emerging studies suggesting that the dedifferentiation or transdifferentiation of beta cells under conditions of prolonged metabolic stress could reduce the beta cell area without causing a substantial loss of endocrine area (reviewed in^25,67^). It is tempting to speculate that a loss, or lower initial proportion, of smaller EOs sitting at an early point in the proposed EO renewal pipeline would put an additional metabolic strain on the residual beta cells within larger EOs, driving pathological outcomes such as beta cell dedifferentiation, transdifferentiation, senescence and beta-cell stress^68,69^. Studies demonstrating that excess BMI in childhood is associated with faster progression towards clinical disease are consistent with this concept^70,71^. Finally, although limited by sample size, the data suggest that most ND donors with AAb (even at young ages) have comparable small EOs to age-matched ND donors. This reduces the possibility that a defect in early development in at-risk individuals accounts for the alterations in EO frequency and density observed in clinical disease. In contrast, those with pathological indicators of T1D progression (presence of IA2A AAb^27^) support our proposal that the small EOs are most susceptible to immune-mediated destruction.

Our collective observations have several potential implications for the understanding of normal pancreas development and T1D pathogenesis. Most notably, the data support the idea that beta cell mass expansion continues within a critical time window postnatally. During this period, a transition occurs from smaller to larger EOs, with a greater proportion of the beta cell mass residing in the medium-large EOs by 7 years of age. In contrast, the very early life period where the greatest pancreas expansion occurs correlates with the time when many beta cells exist as single cells or in small clusters. It is currently unclear whether the rate of insulin secretion from these objects differs from larger EOs at any given glucose concentration. However, the fact that small EOs are preserved after STZ exposure in rodents, which may reflect reduced GLUT2 expression (see above), is consistent with such a possibility. Furthermore, the lack of paracrine signals from neighbouring endocrine cells and the incomplete architecture of the vasculature and innervation will also likely lead to differential responsiveness. One possibility is that these beta cells secrete insulin continuously rather than in response to varying glucose concentrations, thereby acting as a trophic source for pancreatic expansion. Therefore, if these small Ins+ EOs are disrupted (e.g. through priming autoimmunity or via beta cell stressors), their loss will strongly impact both endocrine mass expansion and pancreas growth. This fits with the observed reduction in pancreas weight in individuals with T1D and in those with specific monogenic forms of diabetes caused by enhanced beta cell stress^41,72^.

In summary, we document profound changes in the architecture of the pancreas, particularly in the density and area of EOs in early life. We have confirmed and extended recent 3D studies reporting the presence of single cells and small clusters of beta cells within the pancreas, observing them in donors of all ages without T1D. Importantly, we show that the single cells and small clusters of beta cells are virtually absent in individuals with T1D. We demonstrate that the density of larger EOs is lower in individuals with T1D and that this is most evident in the children who were youngest at clinical diagnosis. We propose that an early and selective elimination of single beta cells and small clusters of beta cells in young individuals may reduce their ability to generate a full complement of endocrine mass during development, resulting in more aggressive clinical disease.

This altered perspective has important implications for current and future treatments and for efforts to implement screening for individuals at-risk of T1D. Our findings imply that a major proportion of beta cells is lost during the islet isolation process. We speculate that the loss of these small EOs during isolation could help explain why whole pancreas transplantation is more successful than isolated islet transplantation, which frequently requires multiple transplants to reverse T1D durably. These findings substantiate a need for early, targeted immunotherapy intervention in Stage 1 in high-risk individuals that could prevent loss, or promote the development of, single beta cells and clusters allowing larger, better protected islets to develop. Furthermore, this study provides support for islet or stem cell-derived islet replacement strategies, especially in individuals diagnosed with T1D at a young age, who may not be able to generate a full complement of endocrine mass; i.e. what was absent initially cannot be protected or recovered (at onset). Finally, this has implications for future beta cell regenerative therapies, which might selectively promote the formation of single beta cells or small clusters that are targeted by active autoimmunity, so a combination with immunotherapeutic agents will be important to protect the newly formed cells.

### Limitations of the study

The present study is limited in several ways, most notably by its cross-sectional nature, precluding interpretations of ‘loss’ or ‘reduction, and the availability of donors from certain age/disease groups. Nonetheless, we report on more donors than any previous study of pancreas morphology, particularly those with T1D. A mixture of post-mortem, organ-donor, and live donor pancreas tissue was included for study, owing to the nature of the biobanks from which tissue was procured. However, we observed no discernible differences in outcomes from these when compared. We also supplemented the work using archival donor tissue immunolabelled for insulin and glucagon, hence we cannot exclude the possibility of other endocrine cell subsets that may exist within the small EOs, beyond alpha and beta cells. This will be the subject of ongoing studies in available tissue. Most of the AAb+ donors available were aged ≥18y (12/15; 80%), and there is limited information available on rates of progression towards T1D in this age group. Finally, we acknowledge that our quantification was undertaken in sections from a mixture of pancreatic regions, and that variation between these is likely.

## STAR METHODS

### RESOURCE AVAILABILITY

#### Lead contact

Requests for further information and resources should be directed to the lead contact, Sarah Richardson

#### Materials availability

Select tissue samples and images from donors evaluated as a part of this study can be requested from nPOD for use in projects approved by the nPOD Tissue Prioritization Committee, as outlined on the nPOD website (www.npod.org), or through approach to the specific biobank curators. This study did not generate new unique reagents.

#### Data and code availability

Spatial imaging data (.qptiff/.svs and .csv), HALO AI classifiers (.onnx), islet and tissue annotation files (.haloannotations) and .R scripts used to perform data pre-processing, analysis and visualisation for this paper will be shared by the lead author upon reasonable request.

### STUDY PARTICIPANT DETAILS

Pancreas tissue from human organ donors with or without T1D was procured for the Network for Pancreatic Organ Donors with Diabetes (nPOD) program at the University of Florida (RRID: SCR_014641, https://www.jdrfnpod.org<u>) and</u> processed in accordance with the established standard operating procedures of the nPOD/OPPC as approved by the University of Florida Institutional Review Board (IRB201600029) with strict adherence to the guidelines of the United Network for Organ Sharing (UNOS) and federal regulations, with informed consent obtained from each donor’s legal representative. The Exeter Archival Diabetes Biobank (EADB) is held with ethical permission from the West of Scotland Research Ethics Committee (ref: 25/WS/0017; IRAS Project ID: 354341). Human fetal tissues were obtained from the Human Developmental Biology Resource (HDBR). These samples were collected with appropriate maternal written informed consent and approval from the Newcastle and North Tyneside (Newcastle University) and London—Fulham (UCL) Research NHS Health Authority Joint Ethics Committees. HDBR is regulated by the UK Human Tissue Authority (HTA; www.hta.gov.uk) and operates in accordance with the relevant HTA Codes of Practice. The DiViD study was approved by The Norwegian Governments Regional Ethics Committee. Written informed consent was obtained from all cases after oral and written information from the diabetologist and the surgeon separately. Three women and three men, three to nine weeks after diagnosis, (aged 24-35 years) participated. Detailed clinical characteristics have been described^73^. Organs for the MRC QUOD Whole Pancreas Biobank were retrieved after informed and written donor family consent in compliance with the UK Human Tissue Act of 2004 under specific ethical approvals by the UK Human Research Authority (05/MRE09/48 and 16NE0230).

### ONLINE METHOD DETAILS

#### Pancreas Tissue

Donor pancreata were obtained from the Exeter Archival Diabetes Biobank (EADB), Human Developmental Biology Resource (HDBR), Network for Pancreatic Organ Donors with Diabetes (nPOD), including small subset of archival samples from Seattle Children Hospital, European nPOD (EUnPOD), and the Diabetes Virus Infection Study (DiViD) biobanks. A total of 334 tissue sections from 220 unique donors were analysed. All donor information and staining performed can be found in **Supplementary Table 1**. Pancreas sections (4 µm) from the Exeter Archival Diabetes Biobank (EADB) and the Human Developmental Biology Resource (HDBR) were labelled for a combination of chromogranin-A (CgA) and cytokeratin-19 (CK19; 10 fetal donors^74^ and 35 neonatal/paediatric donors). An additional donor cohort was obtained from the EADB, nPOD, Seattle and DiViD biobanks in the form of whole slide scans of previously labelled archived tissue, some of which was performed in-house; insulin/glucagon (n=106) and insulin/glucagon/CD45 (n=85).

#### Staining of pancreas sections

Pancreas sections were de-waxed with either 0.5% iodine in xylene or Histo-Clear (National Diagnostics, USA) and rehydrated using decreasing concentrations of ethanols. Heat-induced epitope retrieval in citrate buffer (pH 6.0) was used to unmask antibody-binding epitopes. Sections were blocked in tris-buffered saline (TBS) with 5% normal goat serum before the application of primary antibodies. A dual anti-mouse horseradish peroxidase (HRP) and anti-rabbit alkaline phosphatase (AP) secondary antibody was applied, and a chromogen detection kit was used for secondary antibody detection (HRP-diaminobenzidine (DAB) and AP-Warp Red; Biocare Medical, USA) in slides immunolabelled with either CgA/CK19 or insulin/glucagon. For sections immunolabelled with insulin glucagon and CD45 the chromogen detection reagents DAB, AP-warp red and AP-Ferangi Blue chromogen were used. Sections were mounted using either EcoMount (Biocare Medical, USA) or DPX Mountant for histology (Merck, UK).

### QUANTIFICATION AND STATISTICAL ANALYSIS

#### Imaging and analysis of EO architecture

Labelled sections were digitised at 20x magnification using a PhenoImager™ HT slide scanner (Akoya Biosciences, USA) or an Aperio CS2 slide scanner (Leica Biosystems Imaging, Inc. Whole slide scans were imported into the HALO image analysis platform (Indica Labs, USA). User-created annotations were used to train the DenseNet V2 convoluted neural network to demarcate tissue compartments of interest (connective tissue, acinar, duct, single endocrine cells and those within small clusters and islets). The real-time tuning feature was used to assess classifier performance across iterations. The model was trained until the cross-entropy, a measure of difference between the network classification and user created annotations, reached its lower value and plateaued, as the neural network converged on a solution. Separate DenseNet V2 classifiers were trained for each staining (CgA/CK19, insulin/glucagon & insulin/glucagon/CD45). Each set of classifiers was applied to whole slide scans, to calculate tissue level parameters (total tissue, acinar, endocrine areas) and individual EO parameters (area, diameter, perimeter).

For the insulin/glucagon-labelled sections, the HALO 3.6 area quantification algorithm was applied to calculate the insulin and glucagon positively labelled area within each endocrine object. User-defined RGB values designate staining colours, and an OD threshold sets the intensity for positive staining. Performance of the area quantification algorithm was reviewed on a section-by-section basis, as the optical density varied among historically labelled sections and was optimised if necessary.

Individual EO data were imported into ‘R’ for processing, data visualisation and statistical analysis. Transformation of EO area into the EO bins was performed using the formula:

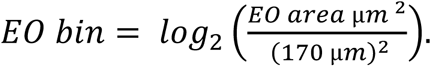

The area of each endocrine object was divided by 170µm^2^, the approximate size of an endocrine cell^17^, to calculate an approximate cell number per EO, followed by log2 transformation. The final EO bin value was rounded down to the nearest integer. This size scale allows for fine detail of the smaller EOs and larger bin sizes for the lower number of larger EOs^17^. Individual EO data and acinar tissue area were passed forward to calculate the EO size profiles, EO density (mm^2^) and % endocrine area/acinar area. EO morphological characteristics (Feret diameters, solidity and circularity^17^) were measured in QuPath v5.1. Data were visualised in *‘R’* (4.1) using *ggplot2*.

#### Statistical analysis

Statistical analysis of data was performed in ‘*R’* (4.1) using the *rstatix* (0.7.2) and *emmeans* (1.11.0) and *car* (v3.1.2) packages. Since the data presented are from a population study where n numbers cannot be controlled experimentally, statistical tests that allow for unbalanced design were performed. The non-parametric Mann-Whitney U test was performed between two groups with a non-normal distribution, followed by the “Benjamini–Hochberg” procedure to calculate adjusted *p* values.

Where three or more groups were compared across the same variable, the Kruskal-Wallis test was performed. *Post hoc* comparisons of all groups were performed using Tukey’s multiple comparison test with the *emmeans* package (v1.11.0) in ‘*R*’.

The type ‘II’ sum of squares ANOVA was performed with the *car* package (v3.1.2), where two or more groups are compared across multiple bin sizes or EO classes. Subsequent *post hoc* testing with Tukey’s multiple comparison test was performed when comparing all selected groups to each other, whereas Dunnett’s test was used for comparing select groups to the ND control group. Specific contrasts computed are indicated in the legend, and all statistical test results can be found in **Supplementary File 1**. Data are presented as mean ± 95% confidence interval (CI). Significance values are denoted as: *, *P* < 0.05; **, *P* < 0.01; ***, *P* < 0.001.

## Supporting information

Main Figures

## ADDITIONAL RESOURCES

Data Portal of the Network for Pancreatic Organ Donors with Diabetes, contains detailed information about each donor tissue used in this study: https://portal.jdrfnpod.org/. Details regarding the EADB Biobank can be found at the NCBI BioSample Resource (BioProject: PRJNA839725), the RRID portal: https://rrid.site/rin/rrids, and on the pancreatlas website https://www.pancreatlas.org/datasets.

### Key resources table

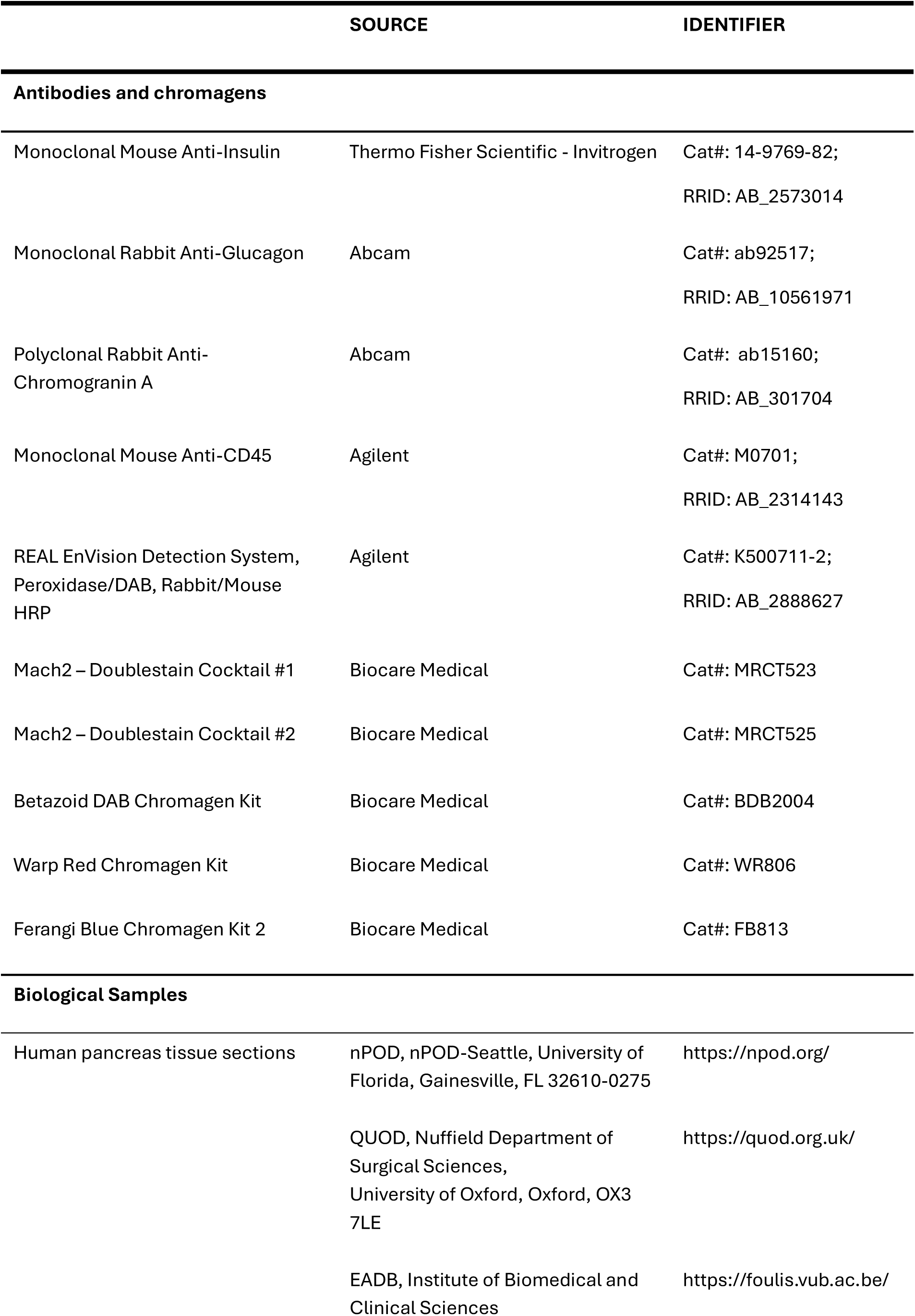
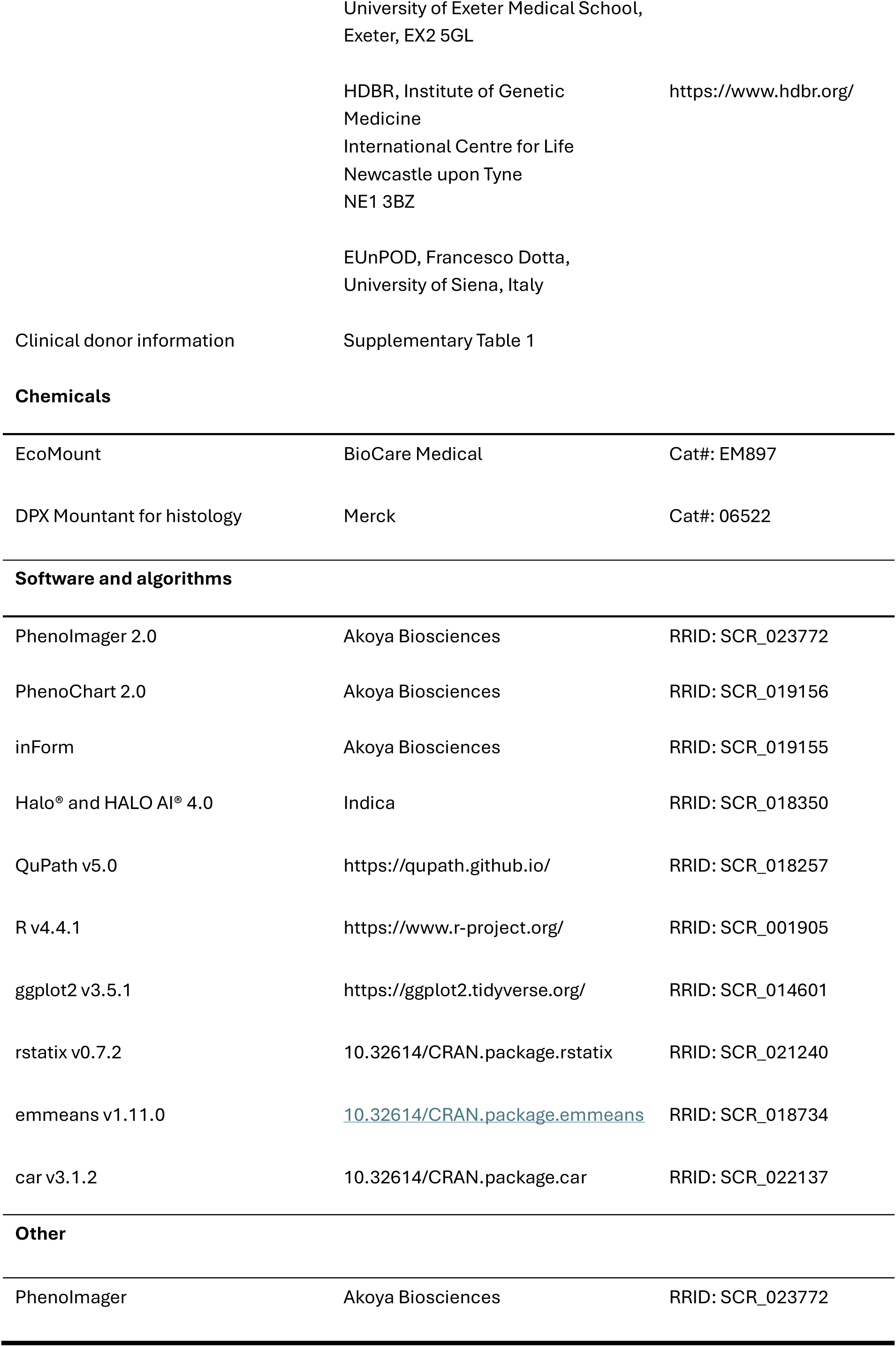

### Supplementary Material

**Supplementary Figure 1 – Quality control checks ensure confounding factors such as immunolabelling marker selection and tissue collection method do not significantly alter EO profiles. (A)** Representative EO shown for calculation of Feret diameters with the minimum and maximum diameter plotted for ND pancreata in bin sizes 0-9. **(B)** Scatter plot and linear regression comparing EO density in serially labelled chromogranin A/cytokeratin-19 (CgA/CK19) and insulin/glucagon (Ins/Gluc) sections. **(C)** Proportion of EO count, proportion of endocrine area and EO density in bins in age matched ND and T1D donors, comparing whether collection method affects EO profile. Data are presented as mean ± 95% CI.

**Supplementary Figure 2 – EO profile shifts towards larger objects with age, with the greatest changes occurring in the first few years post-birth. (A)** Acinar:Endocrine area ratio in pancreata labelled for CgA/CK19 (n=45), grouped by age. **(B)** Proportion of EO count and endocrine area in each bin in ND pancreata labelled for CgA/CK19 and grouped by age. **(C)** Sum of the proportion of EO area in medium sized EOs (bins 4-6) for pancreata labelled for CgA/CK19. **(D)** Proportion of EO count, endocrine area and EO density in each bin for ND pancreata labelled for Ins/Gluc (n=62), grouped by age. **(E)** % endocrine area in each EO bin in ND pancreata labelled for Ins/Gluc, grouped by age. **(F)** Sum of the proportion of EO area in medium sized EOs (bins 4-6) in ND pancreata, grouped by age. Data are presented as mean ± 95% CI. Kruskal-Wallis test followed by Tukey’s *post hoc* test performed to calculate *p* values for **(C)** and **(F)**. *p <0.05, **p <0.01, ***p<0.001.

**Supplementary Figure 3 – Representative spatial plots of the developing pancreas post-birth through to adulthood. (A and B)** Representative spatial plots showing development of endocrine architecture in the neonatal pancreas through to adulthood. Each EO is represented by a single point where its size corresponds to EO bin. The colours of the points designate either size **(A)** or class **(B)**. Each section is labelled with donor ID, age and biobank.

**Supplementary Figure 4 – Beta cells within larger endocrine objects in older individuals are more likely to persist in type 1 diabetes. (A)** Proportion of EO count in each bin, separated by class for ND (n=62) and T1D (n=114) pancreata. **(B)** Schematic for the calculation of circularity and solidity. **(C)** Changes in circularity and solidity in EO bins for ND and T1D pancreata. **(D)** Proportion of EO count in each EO bin, separated by class for ND and T1D (duration ≤2y) pancreata, grouped by age. Type II ANOVA with Tukey’s *post hoc* test was used to calculate *p* values when comparing ND to T1D for each bin. Data are presented as mean ± 95% CI.

**Supplementary Figure 5 – Individuals with IA2A+ autoantibody, or multiple autoantibodies prior to clinical onset of type 1 diabetes, show a loss of small insulin+ only EOs and more small glucagon+ only EOs. (A and B)** Proportion of EO count **(A)** and endocrine area **(B)** for each bin separated by class, comparing age-matched ND, single or double autoantibody positive and T1D pancreata. **(C)** % Insulin or glucagon area in EO bins, comparing age-matched ND, single or double autoantibody positive and T1D pancreata. **(D** and **E)** Proportion of EO count **(D)** and endocrine area **(E)** for each bin separated by class, comparing age-matched ND, single and double autoantibody positive donors, grouped by the presence of IA2A autoantibodies. **(F)** % Insulin or glucagon-labelled area in bins, comparing age-matched ND, single and double autoantibody positive donors grouped by the presence of IA2A autoantibodies. Type II ANOVA with Dunnett’s *post hoc* test comparing autoantibody positive donors to control ND donor pancreata for each bin, was performed to calculate *p* values. Data are presented as mean ± 95% CI.

**Supplementary Figure 6 - Spatial plots of all donors with type 1 diabetes aged ≥13y with a diabetes duration of ≥2y, grouped by endotype. (A and B)** Spatial plots of T1D donors aged ≥13y with a diabetes duration of ≥2y, **(A)** grouped by endotype (T1DE1; diagnosed ≤13y) or T1D endotype 2 (diagnosed ≥13y) **(B)** and ordered by diabetes duration. Each EO is represented by a single point where its size corresponds to its EO bin, and colour denotes its class.

**Supplementary Figure 7 – Characterisation of EO profile, density and % endocrine area in age and duration-matched individuals, diagnosed <13y (T1DE1) or ≥13y (T1DE2), shows a profound absence of small Ins+ EOs across all ages in type 1 diabetes. (A-C)** Proportion of EO count **(A)**, endocrine area **(B)**, and EO density **(C)** for each bin separated by class, in age-matched ND and T1D donors grouped by T1D endotype (T1DE1/2) and diabetes duration (≤2y or ≥2y). **(D)** % Insulin or glucagon area for each bin in ND and T1D pancreata. Data are presented as mean ± 95% CI.

**Supplementary Table 1 – Detailed donor information.**

**Supplementary Table 2 – Changes in the proportion of endocrine area within small (bin 0-3) and large (bin 7-9) EO’s, with age.**

**Supplementary Table 3 – Autoantibody-positive donor clinical information and genetic risk score supplied by the nPOD data portal.** https://portal.jdrfnpod.org/.

Supplementary File 1: Statistical Summary

Supplementary File 2: EXE-T1D consortium members

## Acknowledgements

We thank the organ donors and their families. This research was performed with the support of the Network for Pancreatic Organ donors with Diabetes (nPOD; RRID:SCR_014641), a collaborative type 1 diabetes research project supported by Breakthrough T1D and The Leona M. & Harry B. Helmsley Charitable Trust (Grant#: 3-SRA-2023-1417-S-B). Some images in this manuscript were provided by the Network for Pancreatic Organ Donors with Diabetes (nPOD) Online Pathology database: https://aperioeslide.ahc.ufl.edu. The content and views expressed are the responsibility of the authors and do not necessarily reflect the official view of nPOD. Organ Procurement Organizations (OPO) partnering with nPOD to provide research resources are listed at https://npod.org/for-partners/npod-partners/. We thank Knut Dahl-Jørgensen (Oslo, Norway) for access to the DiViD study donor material, James Shaw (University of Newcastle) for access to pancreas weight from the QUOD Pancreas Study, Marika Bogdani and Gail Deutsch (Seattle Children’s Hospital, USA) for providing donor pancreata from Seattle to the nPOD biobank and Francisco Dotta (Siena, Italy) for access to donors from the EUnPOD study. EUnPOD sample collection is part of the INNODIA project, supported by the Innovative Medicines Initiative 2 (IMI2) - Horizon 2020, grant agreement no. 115797-INNODIA. We also thank Beverley Shields (University of Exeter) for her statistical advice and Myriam Padilla (University of Florida) for her technical support.

## Author Contributions

Conceptualization: S.J.R., K.M.; Supervision: S.J.R., N.G.M.; Resources: S.J.R., N.G.M., J.S., I.K.,; Funding acquisition: S.J.R, N.G.M., EXE-T1D; Data Curation: K.M, T.L.; Formal Analysis: K.M, T.L., C.L.; Investigation: S.J.R., K.M, T.L., C.L.; Methodology: C.S.F., R.W., P.A., P.L., I.B., E.O.; Project Administration: S.J.R, I.K.; Writing: Original draft S.J.R, K.M., T.L., N.G.M.; Writing: Review & Editing: S.J.R, K.M., T.L., N.G.M., C.L., C.S.F., R.W., P.A., P.L., I.B., S.L.H., I.K., M.P., E.O., J.S.

S.J.R is the guarantor of this work and, as such, had full access to all the data in the study and takes responsibility for the integrity of the data and the accuracy of the data analysis.

*EXE-T1D Consortium Contributors are listed in the Supplementary Files.

## Funding

This work was supported by a Steve Morgan Foundation Grand Challenge Senior Research Fellowship awarded to S.J.R (22/0006504), a Strategic Research Agreement award from Breakthrough T1D (formally JDRF), grant reference 2-SRA-2018-474-S-B (S.J.R and N.G.M), a Diabetes UK project grant Reference: 16/0005480 (S.J.R and N.G.M), and The Leona M. and Harry B. Helmsley Charitable Trust grant (G-2103–05059).

This research is supported by the National Institute for Health and Care Research (NIHR) Exeter Biomedical Research Centre (BRC). The views expressed are those of the author(s) and not necessarily those of the NIHR or the Department of Health and Social Care.

This work was conducted in collaboration with the UK T1D-Research Consortium: https://type1diabetesresearch.org.uk/

