## Supplementary figures and images for "Small things matter: Lack of extra-islet beta cells in Type 1 diabetes"

### Main Figures

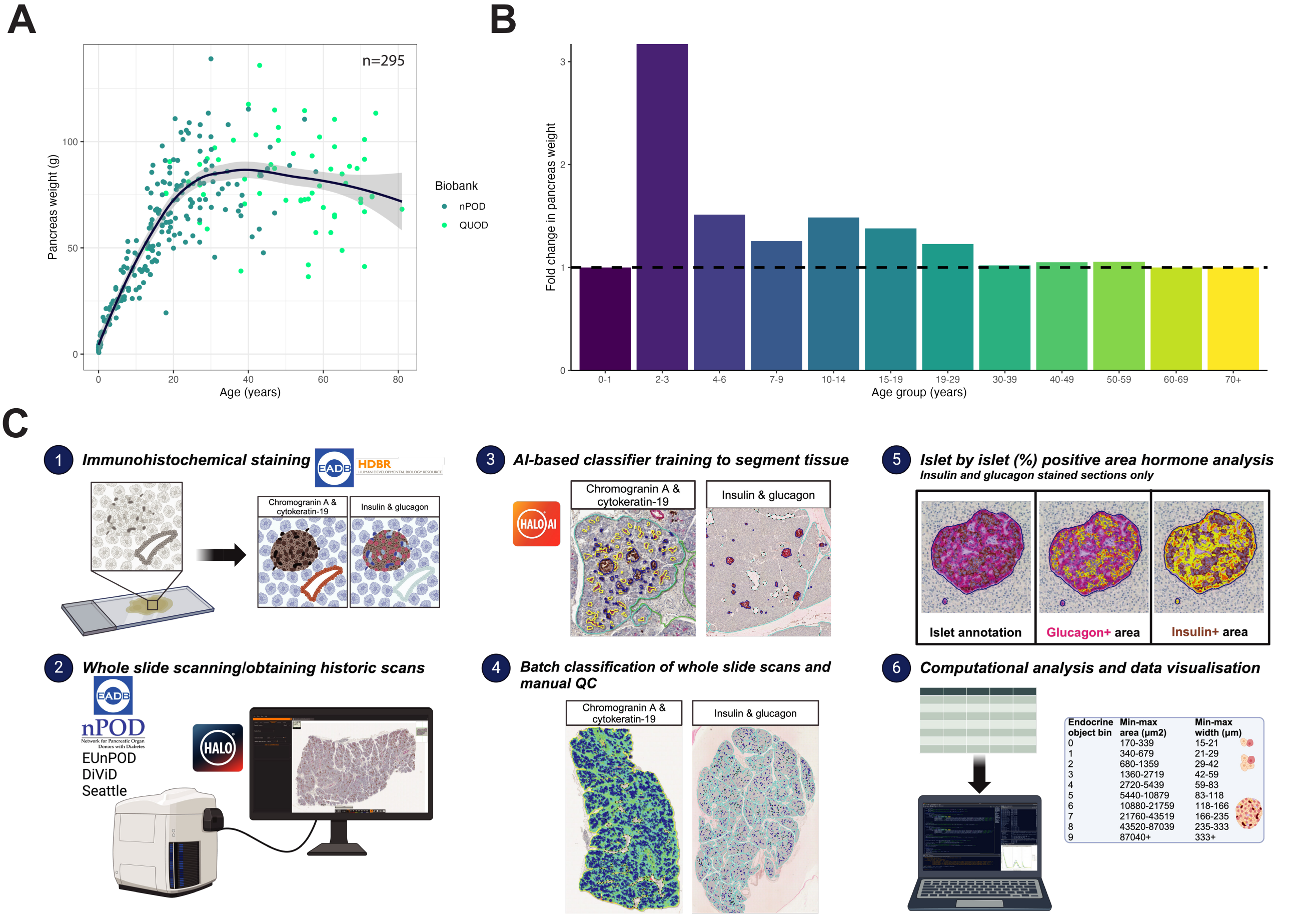

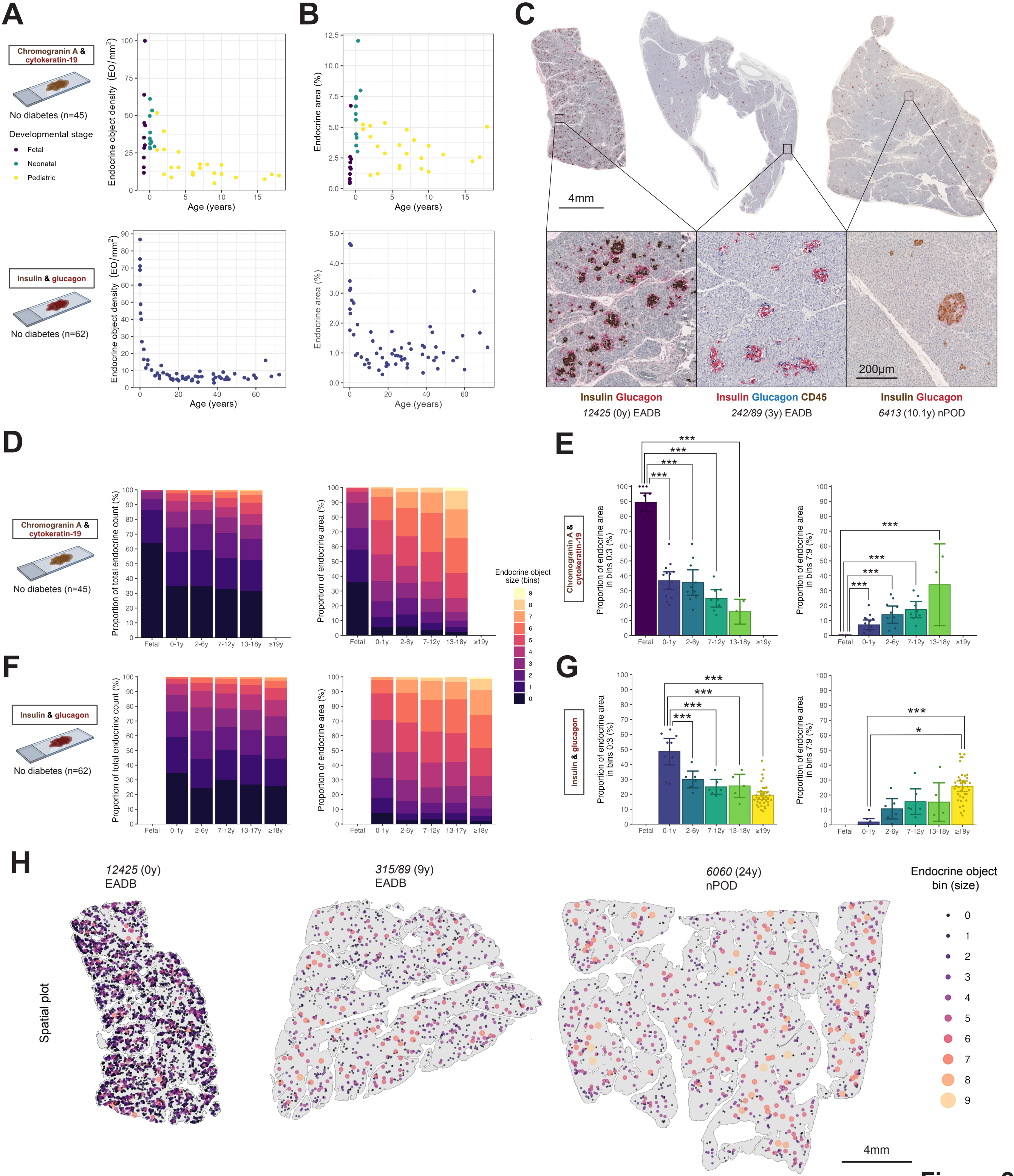

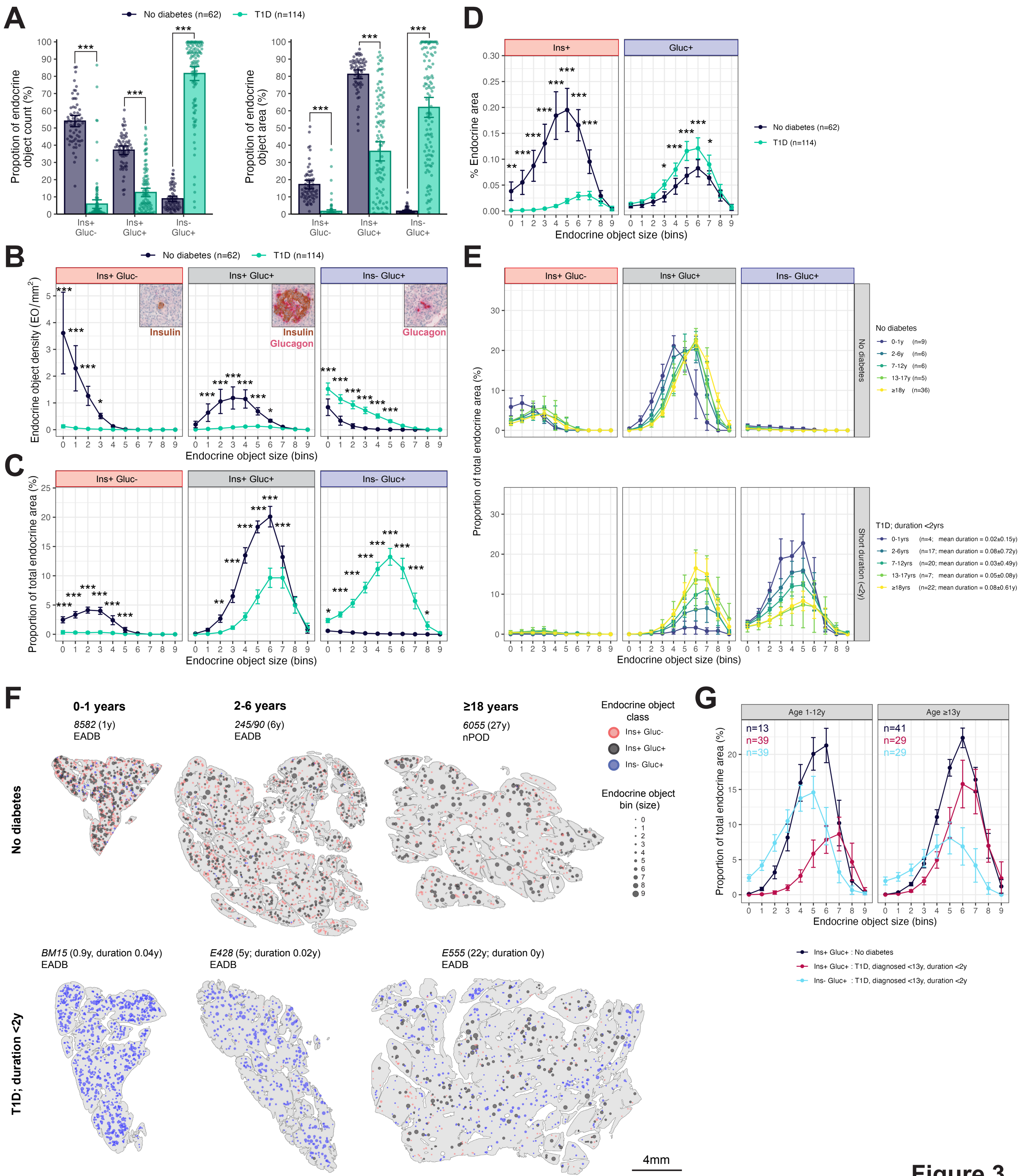

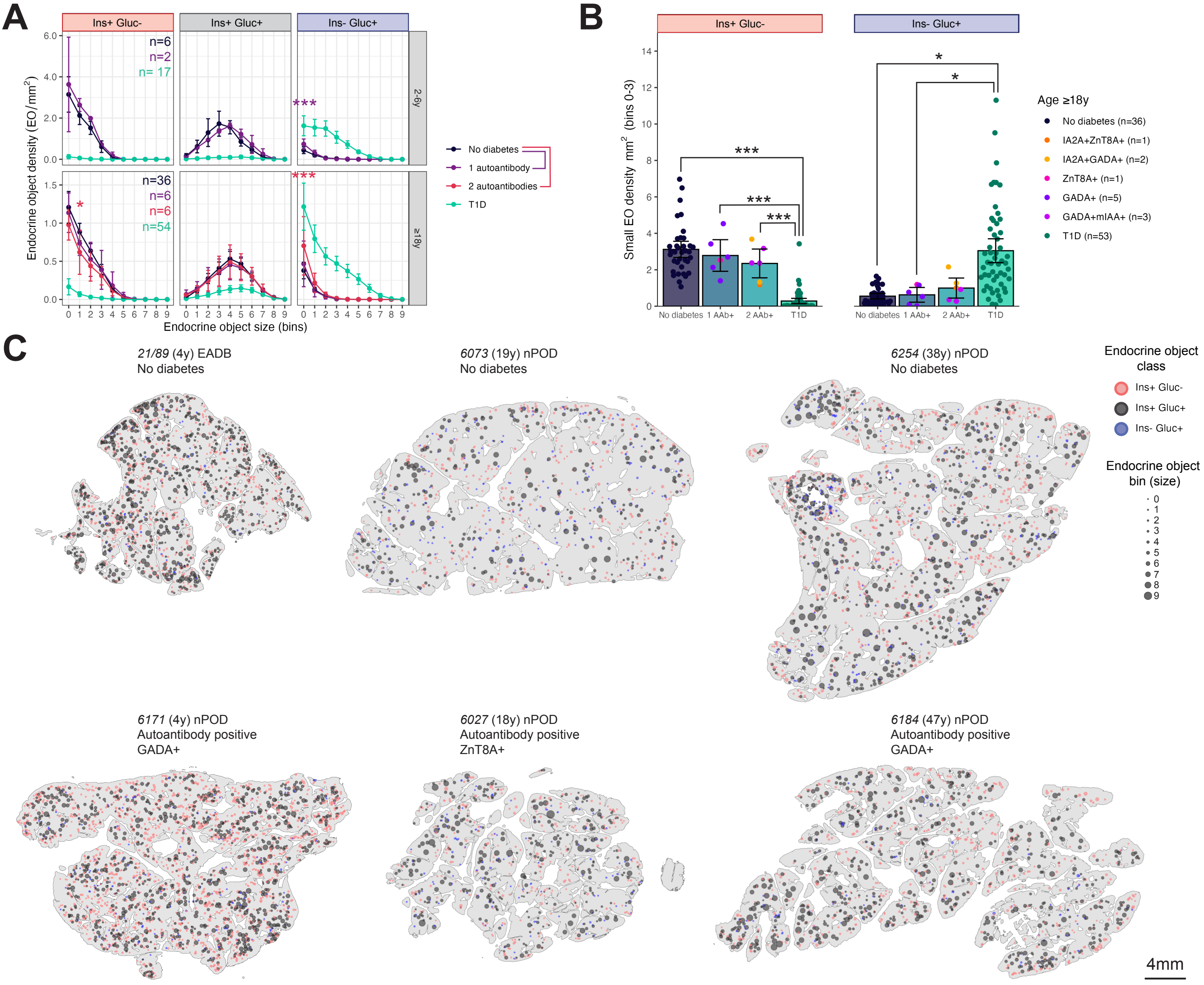

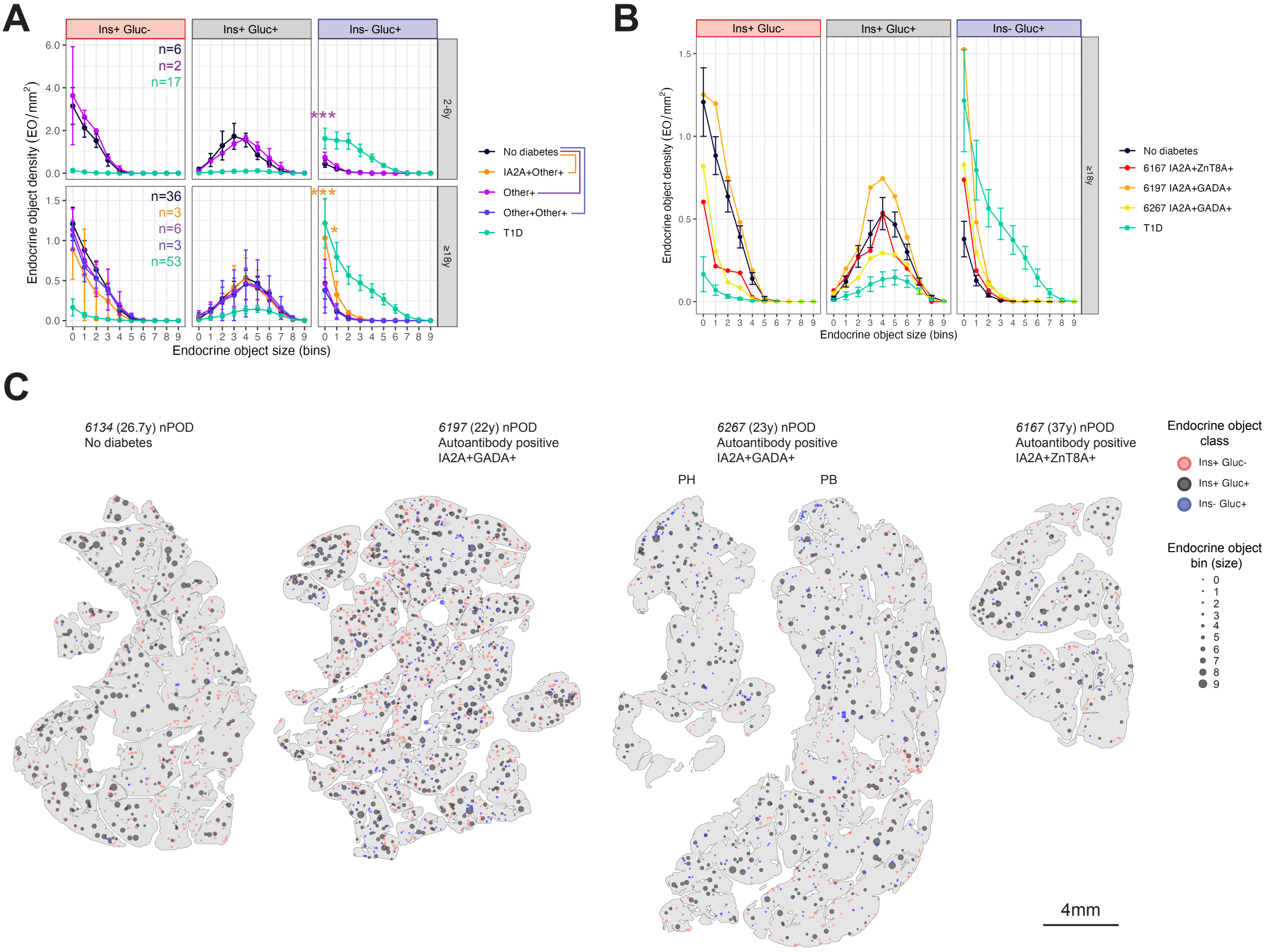

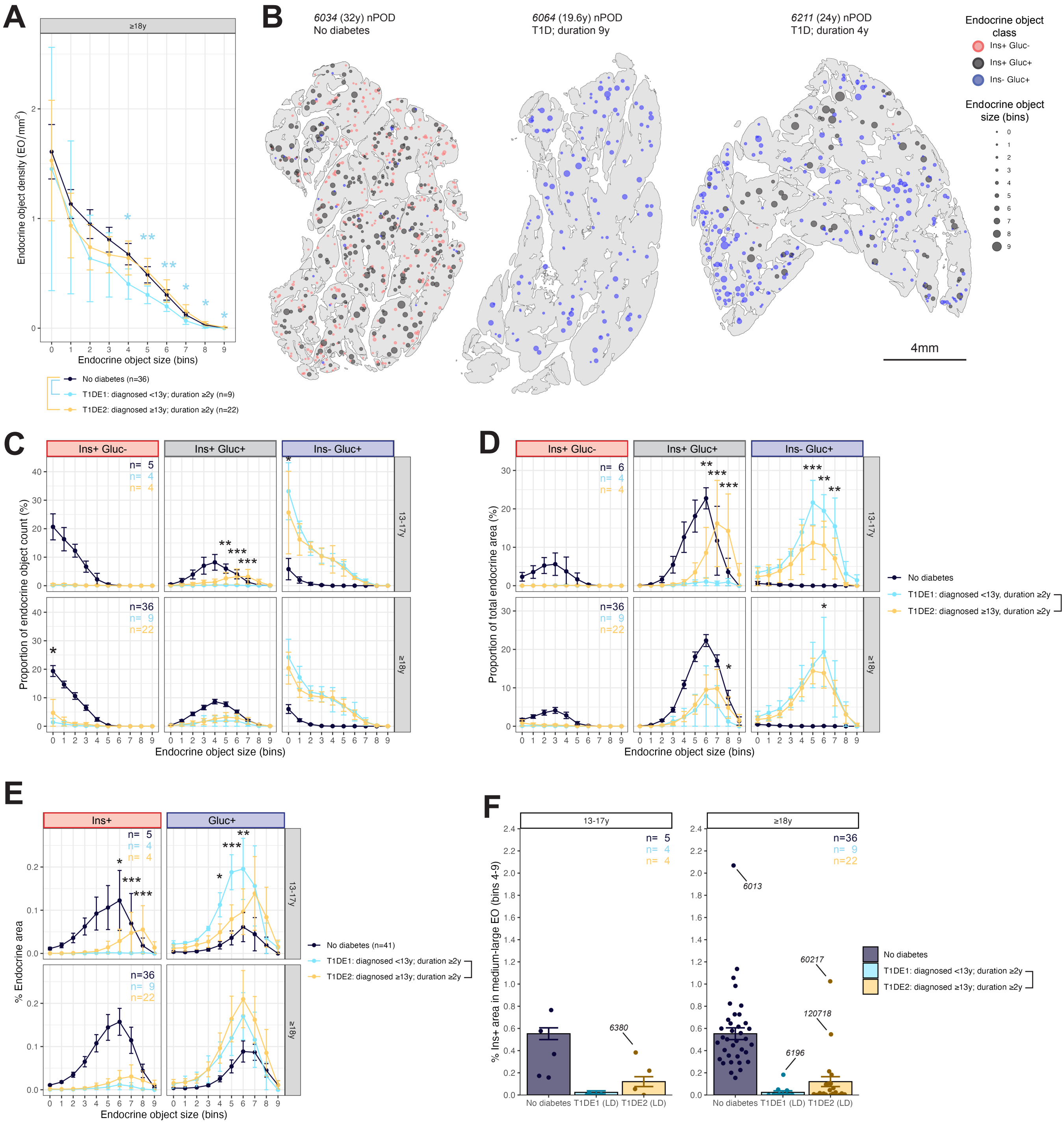
