## Supplementary File 2 for "Small things matter: Lack of extra-islet beta cells in Type 1 diabetes"

**EXE-T1D consortium members:**

Amber M. Luckett^1^, Rebecca A Dobbs^1^, Clara Domingo-Vila^2^, Kathleen M Gillespie^3^, Andrew T Hattersley^1^, Michelle Hudson^1^, Timothy J McDonald^1^, Noel G Morgan^1^, Kathryn Murrall^1^, Sarah J Richardson^1^, Megan E Smithmyer^4^, Cate Speake^4^, Timothy IM Tree^2^, Bart Roep^5^, William A Hagopian^6^, Ben Blaise^7,8^, Iain Yardley^8^, Matthew B Johnson^1^, Richard Oram^1.^

1. Department of Clinical & Biomedical Sciences, University of Exeter Medical School, Exeter, UK

2. Department of Immunobiology, School of Immunology & Microbial Sciences (SIMS), King's College, London, UK

3. Translational Health Sciences, Bristol Medical School, University of Bristol, Southmead Hospital, Bristol, UK

4. Center for Interventional Immunology, Benaroya Research Institute, Seattle, WA, USA

5. Department of Internal Medicine, Leiden University Medical Center, Leiden, the Netherlands

6. Indiana University School of Medicine, Department of Pediatrics, Indianapolis, USA

7. King’s College London, Centre for the Developing Brain, St Thomas’ Hospital, London, United Kingdom

8. Guy's and St Thomas' NHS Foundation Trust, Evelina London Children's Hospital, Department of Paediatric Surgery, London, United Kingdom
